# Down Feather Structure Varies Between Low- and High-Altitude Torrent Ducks (*Merganetta* Armata) in the Andes

**DOI:** 10.1101/207555

**Authors:** Rebecca G. Cheek, Luis Alza, Kevin G. McCracken

## Abstract

Feathers are one of the defining characteristics of birds and serve a critical role in thermal insulation and physical protection against the environment. Feather structure is known to vary among individuals, and it has been suggested that populations exposed to different environmental conditions may exhibit different patterns in feather structure. We examined both down and contour feathers from two populations of male Torrent Ducks (*Merganetta armata*) from Lima, Peru, including one high-altitude population from the Chancay-Huaral River at approximately 3500 meters (m) elevation and one low-altitude population from the Chillón River at approximately 1500 m. Down feather structure differed significantly between the two populations. Ducks from the high-altitude population had longer, denser down compared with low-altitude individuals. Contour feather structure varied greatly among individuals but showed no significant difference between populations. These results suggest that the innermost, insulative layer of plumage (the down), may have developed in response to lower ambient temperatures at high elevations. The lack of observable differences in the contour feathers may be due to the general constraints of the waterproofing capability of this outer plumage layer.

**Resumen:** El plumaje es una característica que define a las aves y cumple roles críticos en el aislamiento térmico y protección física del ambiente. Se sabe que la estructura de las plumas varía ente individuos, y se ha sugerido que poblaciones expuestas a diferentes condiciones ambientales pueden exhibir diferentes patrones en la estructura de las plumas. En este estudio se examinaron tanto el plumón como las plumas de contorno de machos adultos del Pato de los Torrentes (*Merganetta armata*) de dos poblaciones, una en el río Chancay-Huaral a 3,500 msnm y otra en el río Chillón a 1,500 msnm, ubicadas en Lima, Perú. La estructura de los plumones difiere significativamente entre las dos poblaciones. Los patos de la población a grandes elevaciones tienen plumones largos, y densos comparados con los individuos de las partes bajas. La estructura de las plumas de contorno varía ampliamente entre individuos pero no muestra diferencias significativas entre poblaciones. Estos resultados sugieren que las diferencias entre las capas interiores de aislamiento del plumaje (plumón), haberse desarrollado como respuesta en ambientes de bajas temperaturas a grandes elevaciones. En cambio la falta de detectables diferencias en las plumas de contorno puede ser debido a la constante selección en la capacidad impermeable de la capa de plumas exteriores.

## Introduction

Plumage is one of the defining characteristics of birds and serves a critical role in multiple functions including communication, flight, and thermal insulation. Indeed, a reigning theory on the original function of primitive feathers is that they enabled early bird-like dinosaurs to evolve homeothermy (Ostrom 1974; Prum and Brush 2002; Pap et al. 2017), and modern plumage acts as a highly efficient thermal buffer against conductive and convective heat loss both in the air and underwater (Walsberg 1988). All birds shed old and damaged feathers during periodic molts, which is demanding in terms of energy, time, and nutrients so that molting individuals often experience trade-offs between other strenuous periods of the lifecycle such as breeding and migration (Murphy & King 1992). Plumage structure varies between species occupying different habitats (Pap et al. 2017), and is a highly plastic trait that varies between individuals depending on environmental and physiological factors during feather growth (Strochlic & Romero 2008; Butler, Leppert, & Dufty Jr 2010; Moreno-Rueda 2010; Pap et al. 2008, 2013). Therefore, comparing the plumage structure between birds inhabiting different environmental conditions can provide insights into how birds respond to the selection pressures that contribute to this variation.

The body plumage of birds can be broadly divided into two categories: contour, and down. Contour feather structure follows a standard plan of regularly spaced branches (barb) along a central vane (rachis) that has a short basal portion (calamus) imbedded in the skin. Each barb repeats a similar plan with many smaller branches (barbules) densely spaced along either side of the barb. Contour feathers may be further characterized by the exposed, pennaceous (ridged; distal) part of the vane, which aids in water repellency and protection, whereas the plumulaceous (downy; proximal) section provides thermal insulation, recognized through stark differences in barb and barbule texture (Stettenheim 2000; Figure 1). Proportion of plumulaceous barbs, as well as barb and barbule density are thought to determine the amount of air trapped near the skin (Middleton 1986; Butler, Rohwer & Spidel 2008; Broggi et al. 2011, Pap et al. 2017), thereby influencing thermoregulatory capacity (Walsberg 1988). A thicker downy coat composed of longer, denser plumulaceous barbs makes intuitive sense for birds living in colder environments, whereas birds living in hotter environments should have a looser plumage structure to allow them to prevent heat absorption by increasing external surface area of the plumage (Walsberg & King 1978).

**Figure 1.**
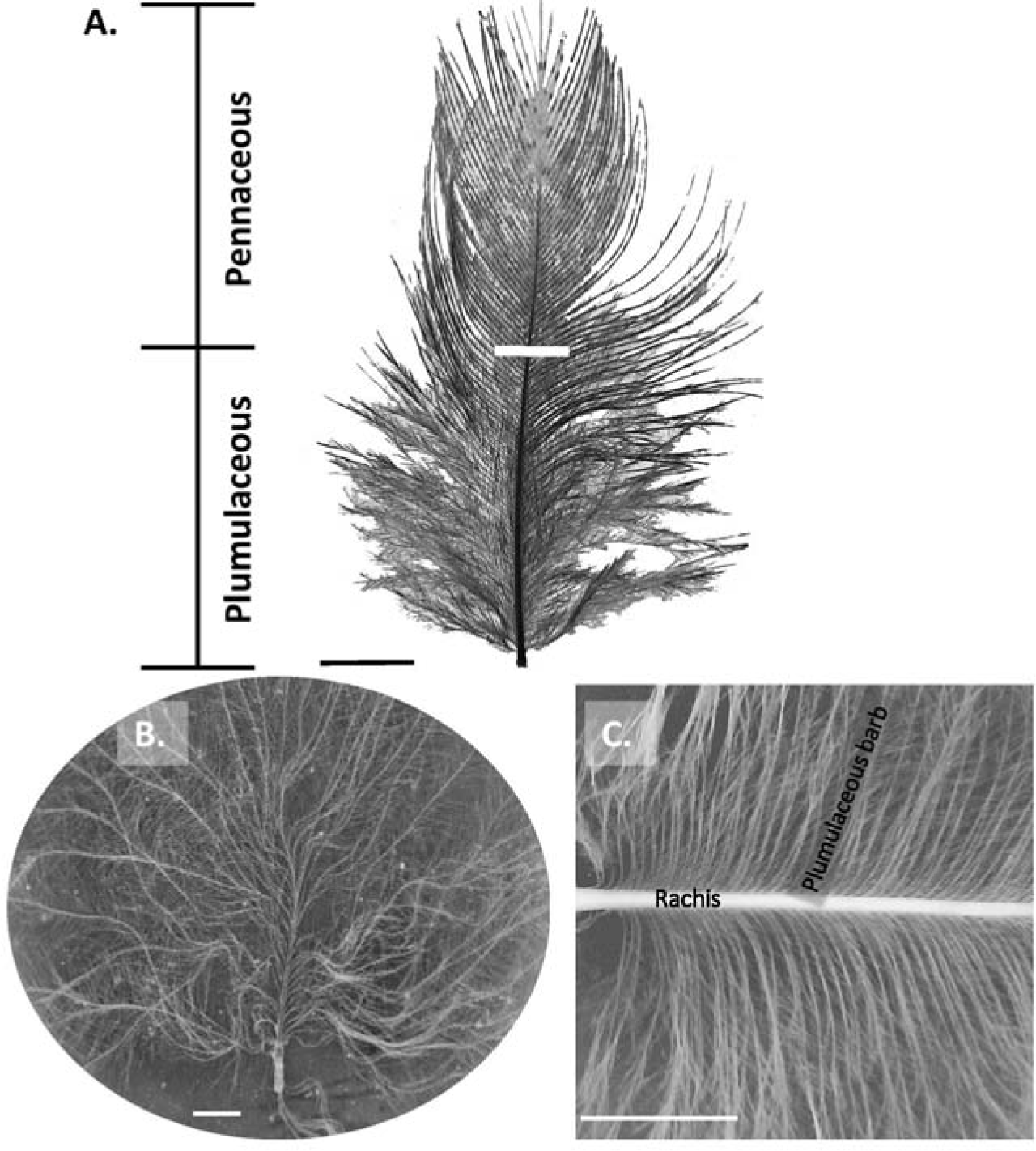
Body feathers of Merganetta armata with plumulaceous and pennaceous sections separated by a white stripe across the rachis. The black lines define the boundary of the distal pennaceous and the proximal plumulaceous portions of the feather (A). Lower figures illustrate M. armata down (B), and a section of plumulaceous vane with the rachis and barb (C). Scale (white bars) for figure (A) and (C) are 0.5 cm, and (B) is 1 mm.

In ducks (Anatidae), contour feathers with diester waxes from the preen gland cover most of the body along discrete tracts and provide an impenetrable waterproof covering over a thick layer of insulating down feathers (Stephenson & Andrews 1997; Stettenheim 2000). As the production of feathers is costly, it would be advantageous for individuals to produce an optimal plumage for the thermal conditions of their given aquatic or terrestrial environment. The energetic costs associated with having suboptimal plumage could be substantial in waterfowl, as thermoregulation can account for 28% of the daily energy expenditure (McKinney & McWilliams 2005). Recent years have seen an increase of studies characterizing interpopulation variation of feather structure due to environmental variation (Middleton 1986; Broggi et al 2011; Gamero et al. 2015; Koskenpato et al. 2016), but few have focused on waterfowl or comparative data describing down feathers among species or populations inhabiting different environments (but see Pap et al. 2017; D’alba et al. 2017).

Torrent Ducks (Merganetta armata) are specialized riverine ducks that inhabit many of the rivers along the Andes from Venezuela to Tierra del Fuego (Fjeldså & Krabbe 1990). This species is characterized as a small bodied (350-550g; Alza et al. 2017) diving duck that forages primarily on aquatic insects by gleaning the surface of submerged boulders (Cerón 2010). M. armata form monogamous pairs, and both sexes cooperate in the defense an approximate 1-2 kilometer stretch of river they inhabit year round (Moffet 1970). M. armata is an ideal organism to study the environmental correlates of feather structure as they occur in elevations that range from 300 to over 4,000 meters (m) (Fjeldså & Krabbe 1990). In Peru, a steep environmental gradient usually consists of an extremely diverse variety of ecological and topographic conditions. On the west slope of the Andes, for example, low-altitude areas along the central coast consist of hot to mild arid deserts interspersed by lush river valleys and lomas (hills) that give way to very cold semi-arid grasslands and glacier-carved montane valleys at higher elevations (Peel, Finlayson & McMahon, 2007; Cheek pers. obs.).

In a recent study of M. armata in the Andes, Gutiérrez-Pinto et al. (2014) found a significant difference in morphological traits, with larger males in low-altitude areas compared to high-altitude areas in Peru. It was speculated that the physiological costs of living in a cold, hypoxic environment is responsible for the observed difference in body mass and skull length along an elevational gradient, but other physiological mechanisms that could allow M. armata to cope in harsh environments (i.e. insulation and daily energy expenditure) have not been fully described. It has further been observed that M. armata in Peru have very little subcutaneous fat (Gutiérrez-Pinto et al., 2014; Cheek pers. obs.), and a concurrent analysis using the same individuals used in this study found no significant difference in resting O^2^ consumption between low- and high-altitude individuals (Ivy pers comm.). Additionally, Dawson et al. (2016) did not find a significant difference in the respiratory (aerobic) capacity in the pectoralis flight muscles of low- and high-altitude M. armata individuals from this study. As flight muscles are credited for a majority of thermogenesis in birds (Butler 1997; Petit & Vézina 2014), these results suggest that these animals do not appear to be altering their metabolic rate to compensate for lower temperatures at higher elevations. Therefore, we predict that M. armata plumage structure will reflect differences due to environmental temperatures associated with low- and high-altitude thermal environments.

In this study, we compared body plumage by examining down and contour feather structure between two populations of M. armata living at two elevational extremes of their distribution characterized by strong differences in environmental temperatures. This allowed us to address the question: Do key structural attributes in M. armata feather insulation differ between populations living in different elevations and temperatures, and do those difference reflect patterns predicted for a diving duck inhabiting cold, hypoxic environments?

## Methods

Twelve adult, male Merganetta armata were collected from two rivers in the Department of Lima, Peru (Appendix); six individuals in the Rio Chancay-Huaral (>3,000 m; Figure 2), and six individuals in the Rio Chillón (<2,000 m; Figure 2), hereafter referred to as the low- and high-altitude populations respectively. The low-altitude study area consists of a mosaic of farmland and thick vegetation along the riverbank, characterized by mean annual temperatures of approximately 20°C and water temperatures of 19°C (Ayala, Ministerio de Agricultura y Riego 2013). The high-altitude study area consists of alpine river valleys with mean annual temperatures of approximately 12°C and water temperatures of 11°C (Vargas, Ministerio de Agricultura y Riego 2015). Though the two populations are geographically isolated in different watersheds, there is moderate levels of gene flow between and across the river systems (Alza et al. unpublished data), indicating that these populations are not genetically structured.

**Figure 2.**
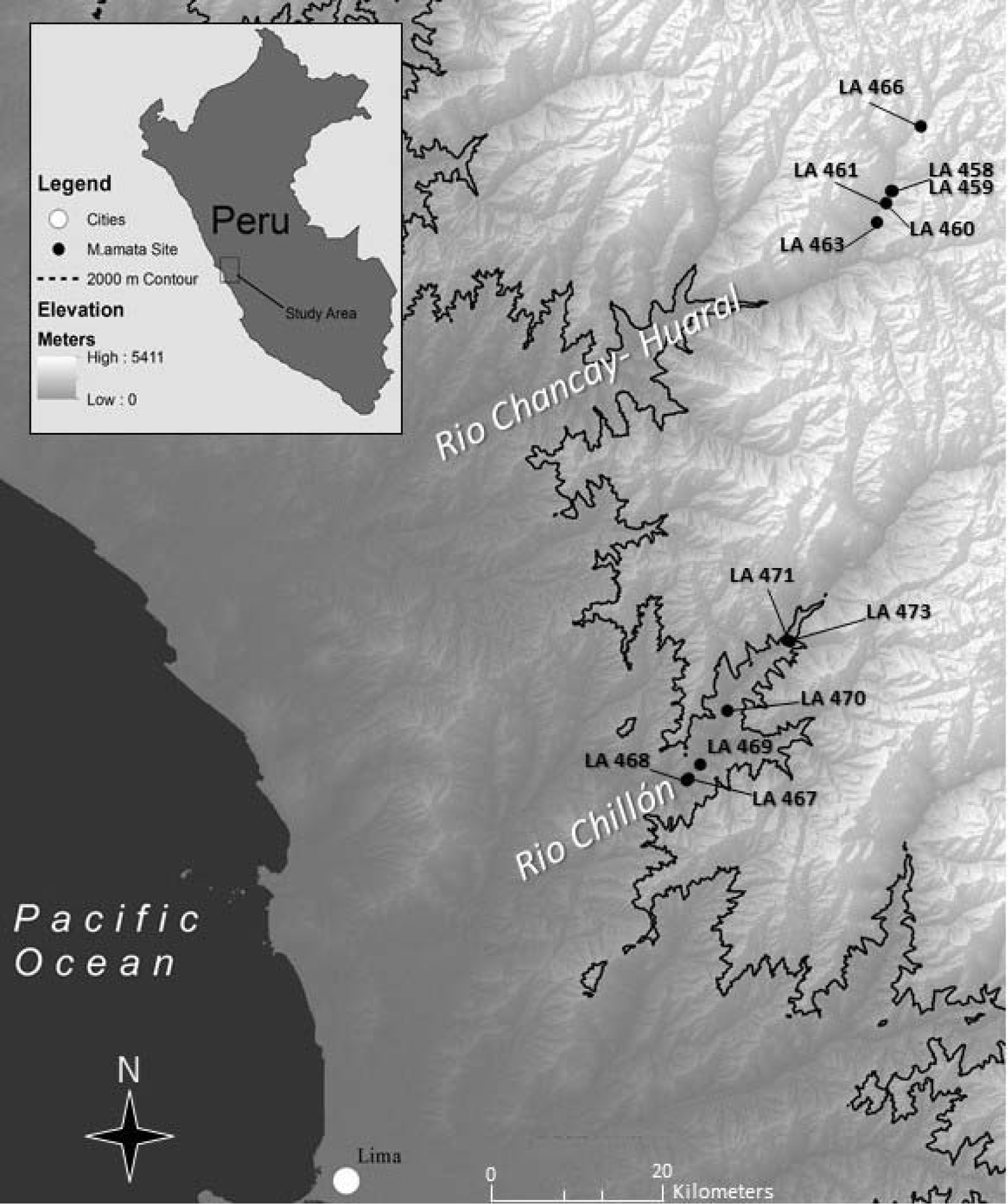
Map of the study areas showing the collection sites of the twelve sampled *Merganetta armata* individuals (Appendix) along the Chillón River (low-altitude population) and Chancay-Huaral River (high-altitude population) in the Department of Lima, Peru.

Ducks from the low and high-altitude populations were euthanized from 7^th^ to 18^th^ August 2015 as part of a concurrent study by the University of Miami and McMaster University (Dawson et al. 2016). There is no reason to expect differences in timing of molt to be a factor, as all animals were sampled within the same two-week time span. Skins were collected from each animal, coated in salt, and stored in a -18°C freezer until subsequent analysis. Feather structure analyses were undertaken in January 2016 in the Centro de Ornitología y Biodiversidad (CORBIDI) laboratory in Lima, Peru. After the skins were thoroughly washed and dried, five down and five contour feathers (minimal sample size for a boxplot comparison, Krzywinski & Altman 2014) were randomly plucked from the upper right pectoral feather tract of each individual. Feathers were plucked and handled with tweezers, and otherwise stored in glassine envelopes. Skins were later deposited in the Ornithology Study Skin Collection at CORBIDI (Appendix).

### Feather Structure

We measured six different traits as described by Middleton (1986) and Broggi (2011) to describe feather structure. Feather length (without calamus) for both down and contour feathers was measured by photographing each feather parallel to a metric ruler. ImageJ software (version 1.50b; Schneider, Rasband, and Eliceiri 2012) was used to calculate feather length with the measuring tool recalibrated for each photo (Figure 1). All photographs and analyses were carried out by the same person (R.C.).

Fine down feather structure was analyzed with the help of a stereoscopic microscope. M. armata down plumage is dense, with barbs regularly spaced along two rachides attached to the calamus. To describe down feather structure, photographs were taken of a single rachis of each feather at 0.8X objective with a camera mounted to the lens. Photographs were analyzed using the ImageJ multi-point count tool to determine total number of barbs along a single rachis for each down feather. Additional feather traits were measured for all contour feathers including: length of plumulaceous portion of each feather, and number of barbs within a 3.5 mm section of the plumulaceous and pennaceous portions of each section (0.8X). Thus, the variables measured were the 1) number of barbs, 2) total length of down feathers; density of barbs from the 3) plumulaceous and 4) pennaceous portions of contour feathers, 5) total length of contour feather, and 6) proportion of plumulaceous barbs with respect to all barbs.

### Statistical Analysis

All data were analyzed in R version 3.3.1 (R Core Team, 2017). All variables except the density of barbs (count data) were normally distributed (Shapiro-Wilk test). To assess the variation in the different feather structure traits by location, we used mixed models controlling for repeated measures (5 feathers per individual). Linear mixed-effects models were used for continuous variables (length and proportion) and generalized linear mixed-effects models for count variables (density of barbs) with a Poisson distribution. Each feather trait was a dependent variable, whereas sampling location (high- and low-altitude populations, fixed effect), and individual (each of the 12 M. armata sampled, random effect) were independent variables.

Confidence intervals for location (fixed effect) variable, were calculated using the R function confint for the non-normally distributed count data, and the function difflsmeans for the continuous data. Finally, we estimated and compared the coefficient of variation for each feather trait by location.

## Results

Down feathers of individuals from high-altitude were, on average, longer and had a greater number of barbs compared to individuals from low-altitude (F_Lenth_=11.815, P_Length_=0.006; F_Barbs_=8.008 P_Barbs_=0.004, df= 10; Figure 3A & 3B). For down length, a 95% confidence interval of the location parameter did not intersect zero CI_Length_ [-3.22, -0.688], and for barb number the parameter did not intersect zero CI_Barbs_ [-0.201, -0.036].

**Figure 3.**
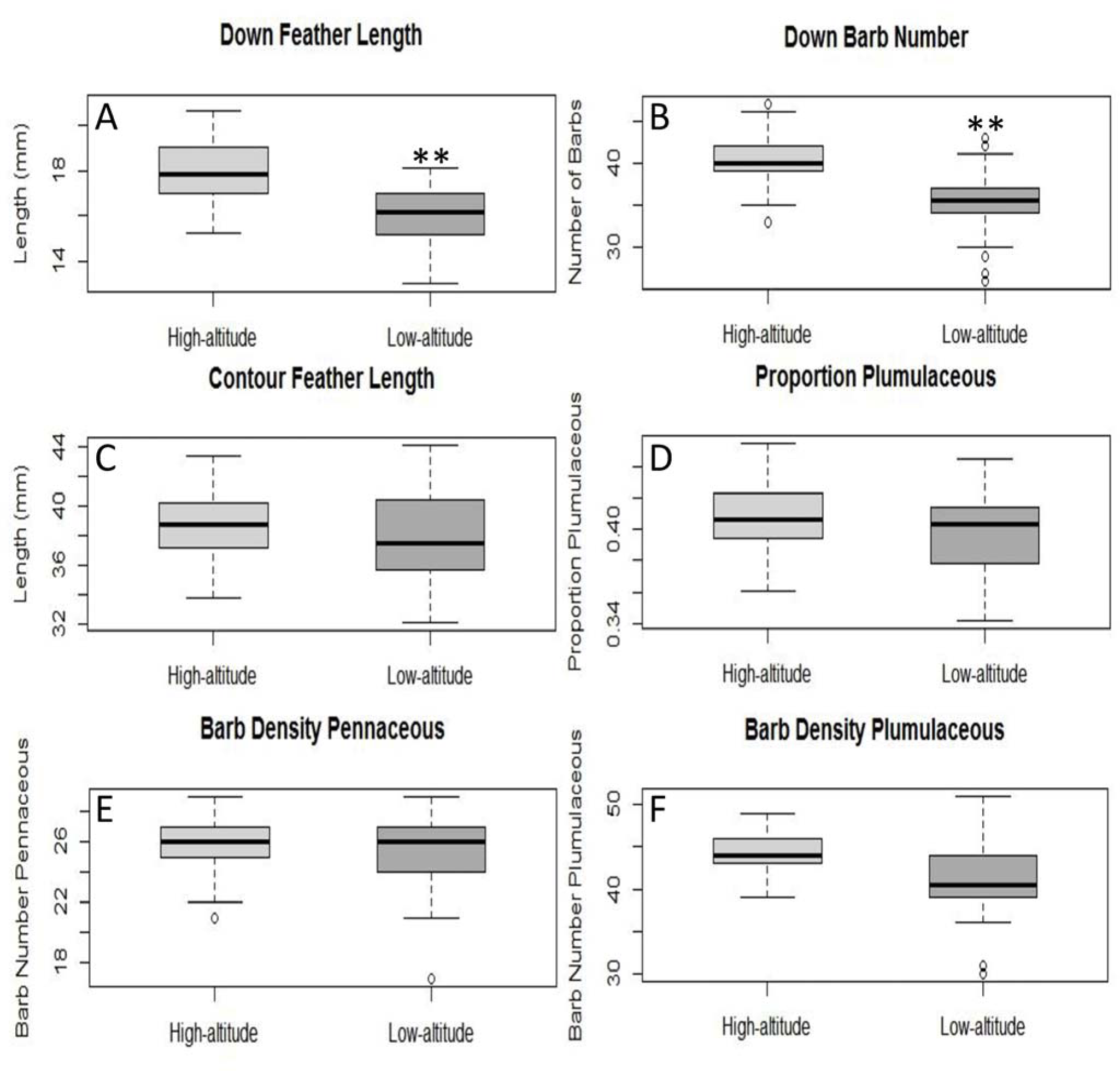
Differences observed in six structural traits of down and contour feathers from the upper right pectoral tracts of twelve adult male Merganetta armata from our high-altitude (light grey), and low-altitude (dark grey) localities in Lima, Peru. Asterisks (*) indicate significant differences between the two populations. Levels of significance: *P < 0.05; **P < 0.01.

Contour feather structure varied greatly between individuals (length) and showed no significant differences between the two populations. Total length of contour feathers did not differ between the populations (F_Length_=0.099, P_Length_= 0.759, df= 10, CI_Length_ [-2.59,3.45], Figure 3C), and the proportion of plumulaceous barbs relative to total number of barbs showed no significant difference between the two populations (F_πPlumulaceous_=1.739, P_πPlumulaceous_= 0.216, df= 10, CI_πPlumulaceous_ [-0.007, 0.03], Figure 3D). Barb number from the plumulaceous section of the feathers did not differ significantly between the two populations (F_BarbsPlumulaceous_=2.364, P_BarbsPlumulaceous_=0.124, df= 10, Figure 3E) as indicated by a 95% confidence interval using of the location parameter CI_BarbsPlumulaceous_ [-0.143,0.021]. The number of barbs from the pennaceous section was highly variable, and the residuals were skewed left by a single noticeable outlier in the low-altitude population. However, when this outlier was removed there were still no observable differences shown between the populations, so the results presented here include all data (F_BarbsPennaceous_=0.1468, P_BarbsPennaceous_= 0.702, df=10, CI_BarbsPennaceous_ [-0.119,0.080]; Figure 3F). Average plumage traits between localities varied more in the low-altitude population compared to the high-altitude individuals sampled (Table 1).

**Table 1.** Coefficient of variation (CV) of feather structure trait averaged between low-altitude and high-altitude populations of adult male Merganetta armata from Lima, Peru.

|  | Down length<br>(mm) | Down barb<br>number | Contour total<br>length (mm) | Contour<br>proportion<br>plumulaceous | Contour barb<br>number-<br>plumulaceous | Contour barb<br>number -<br>pennaceous |
| --- | --- | --- | --- | --- | --- | --- |
| **CV low**<br>**altitude** | **0.0625** | **0.0590** | **0.0713** | 0.0314 | **0.0966** | **0.0613** |
| **CV high**<br>**altitude** | 0.0531 | 0.0286 | 0.0492 | **0.0411** | 0.0369 | 0.0457 |

## Discussion

Down structure differed between low- and high-altitude individuals of adult male Merganetta armata in the west slope of the Andes in Lima, Peru. Merganetta armata sampled from the high-altitude study area (Figure 2) had longer, denser down plumage in the pectoral tract compared to the low-altitude study area (Figure 3). Research has shown that differences in down microstructure are related to differences in insulative properties, as long fibers increase the air-trapping capacity of the feather (D’alba et al. 2017). While our observed differences may appear inconsequential from feather to feather, a difference of 3-4 barbs and 2mm of length in the average down feather across the whole body would be expected to have substantial effects on the overall plumage of the animal. This suggests our observed differences in M. armata down reflect the demands of contrasting environments in the low- and high-altitude temperature regimes of the Peruvian Andes.

The lack of observable differences between the contour feathers of the low- and high-altitude samples could be caused by a diversity of constraints compared with down feathers. First, contour feathers were more variable within individuals (five feathers per individual) than between populations (six individuals per population), which could be due to natural variation among individuals. Second, there is no standard method for quantifying feather structure (Butler et al. 2008), so it is possible that different variables in the contour feathers such as: barb angle (Butler, Rohwer, & Speidel 2008), hue values and infrared spectra (Dove et al. 2007; Gamero et al. 2015), or porosity (a function of barb width and spacing, Rijke 1968, 1970; Rijke & Jesser 2011), are better descriptors of insulation capacity. The methodology applied in this study was used because it was cost effective and easily adapted to an unconventional lab space. Other traits worth investigation are barbule density and feather microstructure (D’alba et al. 2017) of M. armata down and contour feathers. An attempt was made to measure feather barbule density for this study; however, the power of the microscope used was not sufficient to be able to reliably quantify these incredibly minute structures.

Trends in the literature show that analyzing feather structural characteristics is a growing field with a wealth of potential questions that may be answered. Our findings are consistent with a recent phylogenetic review that found no observable differences in the barb density of plumulaceous and pennaceous sections of contour feathers in aquatic birds across environments (Pap et al. 2017). Since M. armata are specialized to a similar habitat type, fast flowing torrential rivers (Johnson 1963), there is no a priori reason to expect contour feathers (i.e., the protective layer of plumage) to differ between low- and high-altitude populations. In contrast to the contour feathers, the insulative down may be more sensitive to environmental temperatures, particularly in aquatic birds. Common garden approaches with passerines have shown that birds from higher latitudes with denser insulative plumage expend less energy on thermoregulation than conspecifics from lower latitudes (Broggi et al. 2005, 2011). Comparative studies between species have also shown that feather characteristics likely reflect the demands of habitat, as aquatic species appear to prioritize waterproofing and plumage cohesion, whereas terrestrial species show greater variation in insulative properties (Pap et al. 2017; D’alba et al. 2017). Future work should build upon this by investigating patterns of plumage variation among populations between habitats, in addition to further comparison of species across habitats.

Our understanding of the importance of down plumage between species is limited as few studies have focused on environmental correlates influencing down structure (but see Williams, Hagelin, and Kooyman 2015; Pap et al. 2017; D’alba et al. 2017). Our findings represent a novel attempt to quantify interpopulation down feather structure between environments along an elevational gradient. Further investigation of down feather structure, particularly in waterfowl, would help to clarify if the observed pattern of longer, denser down plumage in colder environments is observed in other species. Studies investigating feather structure of species across elevational gradients to answer whether plumage is determined through evolutionary processes or a phenotypic plastic response to environmental differences should also be conducted.

## Conclusion

We have shown that high-altitude M. armata have longer, denser down feathers in the pectoral tract compared to low-altitude individuals. Moreover, average plumage traits between localities appeared to vary more in the low-altitude population compared to the high-altitude individuals sampled (Table 1). This could indicate that M. armata plumage is more constrained by selection or developmental plasticity at higher elevations. Further investigation is needed to determine if this pattern is repeated across drainages. These data suggest that these animals are compensating for colder environments by increasing the insulative capacity of their down plumage thereby potentially avoiding further energy expenditures through increased metabolic output for thermogenesis (Dawson et al. 2016; Ivy pers comm). The lack of observable differences in contour feathers may be related to strong constraints of the waterproof capability of this important outer plumage layer in a species that forages underwater. Further investigation is warranted to examine the microstructures of these feathers (D’alba et al. 2017), quantify insulative capacity of the plumage (Walsberg 1988), and determine if these patterns between habitats are repeated across the latitudinal range of M. armata subspecies.

## Acknowledgements

Permits to conduct this research in Peru were provided by SERFOR (Permit: RDG no. 190-2015-SERFOR-DGGSPFFS), and authorization to work in the field was provided by the community of Vichaycocha, and private landowners in Santa Rosa de Quives (Fundo Huanchuy and Hostiliano family). Funding was provided by the University of Alaska Fairbanks Undergraduate Research and Scholarly Activity (URSA) program (awarded to R.C.) and the James A. Kushlan Endowment for Waterbird Biology and Conservation (awarded to K.G.M.) from the University of Miami. We thank G. R. Scott, W. K. Milsom, M. Reichert, B. Chua, J. York, and N. Dawson for logistical support in the field. We thank C. Ivy for physiological data and advice. We are grateful to I. León, L. Alza, C. Corzo, V. Zapata, L. Zapata, B. Alza, R. León, R. Zapata, C. Zapata, and J. Barbarán; Centro de Ornitología y Biodiversidad (T. Valqui, A. Díaz, A. Barboza, P. Venegas, G. Chávez, J. Chávez) for assistance and logistical support in Lima, Peru, and the Dallas World Aquarium (D. Richardson) for laboratory equipment. We would also like to thank D. Kotter for GIS help, and P. Turk for statistical consulting. As well as C. Ghalambor, C. Marshall, J. Harvid, J. Neuwald, and R. Bockrath for comments on earlier drafts. We continuously thank K. Winker for enduring support.

## Appendix

Adult male specimens used in this study with collector’s identifiers (Luis Alza [LA]) catalogue numbers. Collection localities are also included. All voucher specimens are housed in the Centro de Ornitología y Biodiversidad (CORBIDI) ornithology collection, in Lima, Peru.

| Species | Locality | Latitude S | Longitude W | Elevation<br>(meters) | Date collected | Catalogue number |
| --- | --- | --- | --- | --- | --- | --- |
| **High-altitude** | **Rio-Chancay-Huaral** |  |  |  |  |  |
| *Merganetta armata* | Vichaycocha | 11.15983 | 76.62881 | 3680 | 7 Aug 2015 | LA 458 |
| *Merganetta armata* | Vichaycocha | 11.15983 | 76.62881 | 3680 | 7 Aug 2015 | LA 459 |
| *Merganetta armata* | Baños de Collpa | 11.17059 | 76.63445 | 3193 | 8 Aug 2015 | LA 460 |
| *Merganetta armata* | Vichaycocha | 11.15981 | 76.62981 | 3299 | 8 Aug 2015 | LA 461 |
| *Merganetta armata* | Vichaycocha | 11.18813 | 76.64276 | 3000 | 9 Aug 2015 | LA 463 |
| *Merganetta armata* | Vichaycocha | 11.10174 | 76.60341 | 4086 | 10 Aug 2015 | LA 466 |
| **Low-altitude** | **Rio Chillón** |  |  |  |  |  |
| *Merganetta armata* | Santa Rosa de<br>Quives | 11.67533 | 76.80168 | 1092 | 12 Aug 2015 | LA 467 |
| *Merganetta armata* | Santa Rosa de<br>Quives | 11.68945 | 76.81411 | 1034 | 13 Aug 2015 | LA 468 |
| *Merganetta armata* | Santa Rosa de<br>Quives | 11.68801 | 76.81205 | 1040 | 14 Aug 2015 | LA 469 |
| *Merganetta armata* | Fundo Huanchuy | 11.62689 | 76.77708 | 1248 | 15 Aug 2015 | LA 470 |
| *Merganetta armata* | Yaso | 11.56482 | 76.72112 | 1665 | 15 Aug 2015 | LA 471 |
| *Merganetta armata* | Yaso | 11.56362 | 76.72347 | 1615 | 16 Aug 2015 | LA 473 |

